# The ANTsX Ecosystem for Mapping the Mouse Brain

**DOI:** 10.1101/2024.05.01.592056

**Authors:** Nicholas J. Tustison, Min Chen, Fae N. Kronman, Jeffrey T. Duda, Clare Gamlin, Mia G. Tustison, Michael Kunst, Rachel Dalley, Staci Sorenson, Quanxin Wang, Lydia Ng, Yongsoo Kim, James C. Gee

**Author notes:** Corresponding authors: Nicholas J. Tustison, DSc, Department of Radiology and Medical Imaging University of Virginia,; James C. Gee, PhD Department of Radiology University of Pennsylvania.

## Abstract

Precision mapping techniques coupled with high resolution image acquisition of the mouse brain permit the study of the spatial organization of gene expression and their mutual interaction for a comprehensive view of salient structural/functional relationships. Such research is facilitated by standardized anatomical coordinate systems, such as the well-known Allen Common Coordinate Framework (AllenCCFv3), and the ability to spatially map to such standardized spaces. The Advanced Normalization Tools Ecosystem is a comprehensive open-source software toolkit for generalized quantitative imaging with applicability to multiple organ systems, modalities, and animal species. Herein, we illustrate the utility of ANTsX for generating precision spatial mappings of the mouse brain and potential subsequent quantitation. We describe ANTsX-based workflows for mapping domain-specific image data to AllenCCFv3 accounting for common artefacts and other confounds. Novel contributions include ANTsX functionality for velocity flow-based mapping spanning the spatiotemporal domain of a longitudinal trajectory which we apply to the Developmental Common Coordinate Framework. Additionally, we present an automated structural morphological pipeline for determining volumetric and cortical thickness measurements analogous to the well-utilized ANTsX pipeline for human neuroanatomical structural morphology which illustrates a general open-source framework for tailored brain parcellations.

## 1 Introduction

Over the past two decades there have been significant advancements in mesoscopic analysis of the mouse brain. It is currently possible to track single cell neurons in mouse brains,^1^ observe whole brain developmental changes on a cellular level,^2^ associate brain regions and tissues with their genetic composition,^3^ and locally characterize neural connectivity.^4^ Much of this scientific achievement has been made possible due to breakthroughs in high resolution imaging techniques that permit submicron, 3-D imaging of whole mouse brains. Associated research techniques such as micro-optical sectioning tomography,^6^ tissue clearing,^1,7^ spatial transcriptomics^9^ are all well-utilized in the course of scientific investigations of mesoscale relationships in the mouse brain.

An important component of this research is the ability to map the various image data to anatomical reference frames^11^ for inferring spatial relationships between structures, cells, and genetics. This has motivated the development of detailed structural image atlases of the mouse brain. Notable examples include the Allen Brain Atlas and Common Coordinate Frameworks (AllenCCFv3),^13^ the Waxholm Space,^14^ and more recently, the Developmental Common Coordinate Framework (DevCCF).^15^ Despite the significance of these contributions, challenges still exist in large part due to the wide heterogeneity in associated study-specific image data. For example, variance in the acquisition methods can introduce artifacts such as tissue distortion, holes, bubbles, folding, tears, and missing slices. These complicate assumed correspondence for conventional spatial mapping approaches.

### 1.1 Mouse-specific brain mapping software

To address such challenges, several software packages have been developed over the years comprising solutions of varying comprehensibility, sophistication, and availability. An early contribution to the community was the Rapid Automatic Tissue Segmentation (RATS) package^16^ for brain extraction. More recently, several publicly available packages comprise well-established package dependencies originally developed on human brain data. SPMMouse,^17^ for example, is based on the well-known Statistical Parametric Mapping (SPM) Matlab-based toolset.^18^ The automated mouse atlas propagation (aMAP) tool is largely a front-end for the NiftyReg image registration package^19^ applied to mouse data which is currently available as a Python module.^20^ NiftyReg is also used by the Atlas-based Imaging Data Analysis (AIDA) MRI pipeline^21^ as well as the Multi Atlas Segmentation and Morphometric Analysis Toolkit (MASMAT). Whereas the former also incorporates the FMRIB Software Library (FSL)^22^ for brain extraction and DSIStudio^23^ for DTI processing, the latter uses NiftySeg and multi-consensus labeling tools^24^ for brain extraction and parcellation. In addition, MASMAT incorporates N4 bias field correction^25^ from the Advanced Normalization Tools Ecosystem (ANTsX)^26^ as do the packages Multi-modal Image Registration And Connectivity anaLysis (MIRACL),^27^ Sammba-MRI,^28^ and Small Animal Magnetic Resonance Imaging (SAMRI).^29^ However, whereas Saamba-MRI uses AFNI^30^ for image registration; MIRACL, SAMRI, SAMBA,^31^ and BrainsMapi^32^ all use ANTsX registration tools. Other packages use landmark-based approaches to image registration including SMART—^33^ an R package for semi-automated landmark-based registration and segmentation of mouse brain based on WholeBrain.^34^ Relatedly, FriendlyClearMap^35^ and mBrainAligner^36^ are both landmark-based approaches to mapping of the mouse brain. Whereas the former employs Elastix^37^ functionality, the latter is based on developed methodology referred to as *coherent landmark mapping*. Finally, the widespread adoption of deep learning techniques has also influenced development in mouse brain imaging methodologies. For example, if tissue deformations are not considered problematic for a particular dataset, DeepSlice can be used to determine affine mappings^38^ with the optimal computational efficiency associated with neural networks.

### 1.2 The ANTsX Ecosystem for mouse brain mapping

As noted previously, many of the existing packages designed for processing mouse brain image data use ANTsX tools for core processing steps in various workflows, particularly its pairwise, intensity-based image registration capabilities and bias field correction. Historically, ANTsX development is originally based on fundamental approaches to image mapping,^39–41^ particularly in the human brain, which has resulted in core contributions to the field such as the well-known Symmetric Normalization (SyN) algorithm.^42^ Since its development, various independent platforms have been used to evaluate ANTsX image registration capabilities in the context of different application foci which include multi-site brain MRI data,^43^ pulmonary CT data,^44^ and most recently, multi-modal brain registration in the presence of tumors.^45^

Apart from its registration capabilities, ANTsX comprises additional functionality such as template generation,^46^ intensity-based segmentation,^47^ preprocessing,^25,48^ deep learning networks,^26^ and other miscelleneous utilities (see Table 1). The comprehensive use of the toolkit has demonstrated superb performance in multiple application areas (e.g., consensus labeling,^49^ brain tumor segmentation,^50^ and cardiac motion estimation^51^). Importantly, ANTs is built on the Insight Toolkit (ITK)^52^ deriving benefit from the open-source community of scientists and programmers as well as providing an important resource for algorithmic development, evaluation, and improvement. We use this functionality to demonstrate recently developed frameworks for mapping fluorescence micro-optical sectioning tomography (fMOST) and multiplexed error-robust fluorescence in situ hybridization (MERFISH) image data to the AllenCCFv3 atlas space. In addition to standard preprocessing steps (e.g., bias correction), additional considerations are accommodated within the ANTsX ecosystem, such as section reconstruction and landmark-based alignment with corresponding processing scripts available at https://github.com/dontminchenit/CCFAlignmentToolkit.

**Table 1:** Sampling of ANTsX functionality.

| *ANTsPy: Preprocessing* |  |
| --- | --- |
| bias field correction | `n4_bias_field_correction(...)` |
| image denoising | `denoise_image(...)` |
| *ANTsPy: Registration* |  |
| image registration | `registration(...)` |
| image transformation | `apply_transforms(...)` |
| template generation | `build_template(...)` |
| landmark registration | `fit_transform_to_paired_points(...)` |
| time-varying landmark reg. | `fit_time_varying_transform_to_point_sets(...)` |
| integrate velocity field | `integrate_velocity_field(...)` |
| invert displacement field | `invert_displacement_field(...)` |
| *ANTsPy: Segmentation* |  |
| MRF-based segmentation | `atropos(...)` |
| Joint label fusion | `joint_label_fusion(...)` |
| diffeomorphic thickness | `kelly_kapowski(...)` |
| *ANTsPy: Miscellaneous* |  |
| Regional intensity statistics | `label_stats(...)` |
| Regional shape measures | `label_geometry_measures(...)` |
| B-spline approximation | `fit_bspline_object_to_scattered_data(...)` |
| Visualize images and overlays | `plot(...)` |
| *ANTsPyNet: Mouse-specific* |  |
| brain extraction | `mouse_brain_extraction(...modality="t2"...)`<br>`mouse_brain_extraction(...modality="ex5"...)` |
| brain parcellation | `mouse_brain_parcellation(...)` |
| cortical thickness | `mouse_cortical_thickness(...)` |
| super resolution | `mouse_histology_super_resolution(...)` |

ANTsX provides state-of-the-art functionality for processing biomedical image data. Such tools, including deep learning networks, support a variety of mapping-related tasks. A more comprehensive listing of ANTsX tools with self-contained R and Python examples is provided as a gist page on GitHub (https://tinyurl.com/antsxtutorial).

### 1.3 ANTsX-based open-source contributions

Consistent with previous ANTsX development, the newly introduced capabilities introduced below are available through ANTsX (specifically, via R and Python ANTsX packages), and illustrated through self-contained examples in the ANTsX tutorial (https://tinyurl.com/ antsxtutorial) with a dedicated GitHub repository specific to this work (https://github. com/ntustison/ANTsXMouseBrainMapping).

#### 1.3.1 The DevCCF velocity flow model

Recently, the Developmental Common Coordinate Framework (DevCCF) was introduced to the mouse brain research community as a public resource^15^ comprising symmetric atlases of multimodal image data and anatomical segmentations defined by developmental ontology. These templates sample the mouse embryonic days (E) 11.5, E13.5, E15.5, E18.5 and postnatal day (P) 4, P14, and P56. Modalities include light sheet flourescence miscroscopy (LSFM) and at least four MRI contrasts per developmental stage. Anatomical parcellations are also available for each time point and were generated from ANTsX-based mappings of gene expression and other cell type data. Additionally, the P56 template was integrated with the AllenCCFv3 to further enhance the practical utility of the DevCCF. These processes, specifically template generation and multi-modal image mapping, were performed using ANTsX functionality in the presence of image mapping difficulties such as missing data and tissue distortion.^15^

Given the temporal gaps in the discrete set of developmental atlases, we also provide an open-source framework for inferring correspondence within the temporally continuous domain sampled by the existing set of embryonic and postnatal atlases of the DevCCF. This recently developed functionality permits the generation of a diffeomorphic velocity flow transformation model,^53^ influenced by previous work.^54^ The resulting time-parameterized velocity field spans the stages of the DevCCF where mappings between any two continuous time points within the span bounded by the E11.5 and P56 atlases is determined by integration of the optimized velocity field.

#### 1.3.2 Structural morphology and cortical thickness in the mouse brain

One of the most frequently utilized pipelines in the ANTsX toolkit is that of estimating cortical thickness maps in the human brain. Beginning with the Diffeomorphic Registration-based Cortical Thickness (DiReCT) algorithm,^55^ this was later expanded to include a complete processing framework for human brain cortical thickness estimation for both cross-sectional^56^ and longitudinal^57^ data using T1-weighted MRI. These pipelines were later significantly refactored using deep learning innovations.^26^

In contrast to the pipeline development in human data,^26^ no current ANTsX tools exist to create adequate training data for the mouse brain. In addition, mouse brain data acquisition often has unique issues, such as lower data quality or sampling anisotropy which limits its applicability to high resolution resources (e.g., AllenCCFv3, DevCCF), specifically with respect to the corresponding granular brain parcellations derived from numerous hours of expert annotation leveraging multimodal imaging resources.

Herein, we introduce a mouse brain cortical thickness pipeline for T2-weighted (T2-w) MRI comprising two novel deep learning components: two-shot learning brain extraction from data augmentation of two ANTsX templates generated from two open datasets^58,59^ and single-shot brain parcellation derived from the AllenCCFv3 labelings mapped to the corresponding DevCCF P56 T2-w component. Although we anticipate that this cortical thickness pipeline will be beneficial to the research community, this work demonstrates more generally how one can leverage ANTsX tools for developing tailored brain parcellation schemes using these publicly available resources. Evaluation is performed on an independent open dataset^60^ comprising longitudinal acquisitions of multiple specimens.

## 2 Results

### 2.1 AllenCCFv3 brain image mapping

#### 2.1.1 Mapping fluorescence micro-optical sectioning tomography data

##### Overview

A framework for mapping fluorescence micro-optical sectioning tomography (fMOST) mouse brain images into the AllenCCFv3 was developed (see Figure 1(a)). An intensity- and shape-based average fMOST atlas serves as an intermediate registration target for mapping fMOST images from individual specimens into the AllenCCFv3. Preprocessing steps include downsampling to match the 25*µm* isotropic AllenCCFv3, acquisition-based stripe artifact removal, and inhomogeneity correction.^25^ Preprocessing also includes a single annotation-driven registration to establish a canonical mapping between the fMOST atlas and the AllenCCFv3. This step allows us to align expert determined landmarks to accurately map structures with large morphological differences between the modalities, which are difficult to address using standard approaches. Once this canonical mapping is established, standard intensity-based registration is used to align each new fMOST image to the fMOST specific atlas. This mapping is concatenated with the canonical fMOST atlas-to-AllenCCFv3 mapping to further map each individual brain into the latter without the need to generate additional landmarks. Transformations learned through this mapping can be applied to single neuron reconstructions from the fMOST images to evaluate neuronal distributions across different specimens into the AllenCCFv3 for the purpose of cell census analyses.

**Figure 1:**
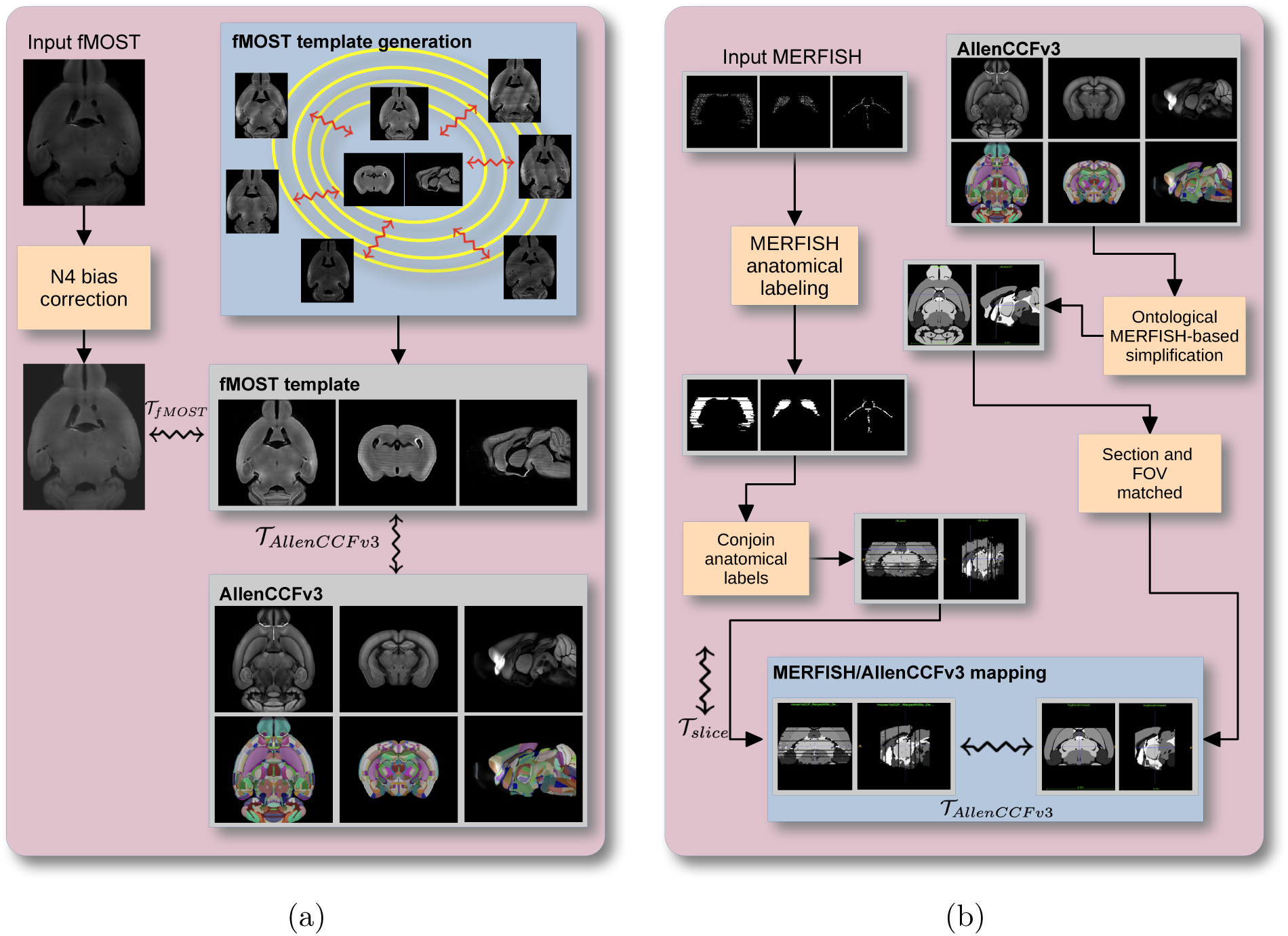
Diagrammatic illustration of the two ANTsX-based pipelines for mapping (a) fMOST and (b) MERFISH data into the space of AllenCCFv3. Each generates the requisite transforms, *T*, to map individual images.

##### Data

The high-throughput and high-resolution fluorescence micro-optical sectioning tomography (fMOST)^61,62^ platform was used to image 55 mouse brains containing gene-defined neuron populations, with sparse transgenic expression.^63,64^ In short, the fMOST imaging platform results in 3-D images with voxel sizes of 0.35 × 0.35 × 1.0*µm*^3^ and is a two-channel imaging system where the green channel displays the green fluorescent protein (GFP) labeled neuron morphology and the red channel is used to visualize the counterstained propidium iodide cytoarchitecture. The spatial normalizations described in this work were performed using the red channel, which offered higher tissue contrast for alignment, although other approaches are possible including multi-channel registration.

##### Evaluation

Evaluation of the canonical fMOST atlas to Allen CCFv3 mapping was performed via quantitative comparison at each step of the registration and qualitative assessment of structural correspondence after alignment by an expert anatomist. Dice values were generated for the following structures: whole brain, 0.99; fimbria, 0.91; habenular commissure, 0.63; posterior choroid plexus, 0.93; anterior choroid plexus, 0.96; optic chiasm, 0.77; caudate putamen, 0.97. Similar qualitative assessment was performed for each fMOST specimen including the corresponding neuron reconstruction data.

#### 2.1.2 Mapping multiplexed error-robust fluorescence in situ hybridization (MERFISH) data

##### Overview

The unique aspects of mapping multiplexed error-robust fluorescence in situ hybridization (MERFISH) spatial transcriptomic data onto AllenCCFv3^65^ required the development of a separate ANTsX-based pipeline (see Figure 1(b)). Mappings are performed by matching gene expression derived region labels from the MERFISH data to corresponding anatomical parcellations of the AllenCCFv3. The pipeline consists of MERFISH data specific preprocessing which includes section reconstruction, mapping corresponding anatomical labels between AllenCCFv3 and the spatial transcriptomic maps of the MERFISH data, and matching MERFISH sections to the atlas space. Following pre-processing, two main alignment steps were performed: 1) 3-D global affine mapping and section matching of the AllenCCFv3 into the MERFISH data and 2) 2D global and deformable mapping between each MERFISH section and matched AllenCCFv3 section. Mappings learned via each step in the pipeline are preserved and concatenated to provide point-to-point correspondence between the original MERFISH data and AllenCCFv3, thus allowing individual gene expressions to be transferred into the AllenCCFv3.

##### Data

MERFISH mouse brain data was acquired using a previously detailed procedure.^65^ Briefly, a brain of C57BL/6 mouse was dissected according to standard procedures and placed into an optimal cutting temperature (OCT) compound (Sakura FineTek 4583) in which it was stored at −80°C. The fresh frozen brain was sectioned at 10*µm* on Leica 3050 S cryostats at intervals of 200*µm* to evenly cover the brain. A set of 500 genes were imaged that had been carefully chosen to distinguish the ~ 5200 clusters of our existing RNAseq taxonomy. For staining the tissue with MERFISH probes, a modified version of instructions provided by the manufacturer was used.^65^ Raw MERSCOPE data were decoded using Vizgen software (v231). Cell segmentation was performed.^66^ In brief, cells were segmented based on DAPI and PolyT staining using Cellpose.^67^ Segmentation was performed on a median z-plane (fourth out of seven) and cell borders were propagated to z-planes above and below. To assign cluster identity to each cell in the MERFISH dataset, we mapped the MERFISH cells to the scRNA-seq reference taxonomy.

##### Evaluation

Alignment of the MERFISH data into the AllenCCFv3 was qualitatively assessed by an expert anatomist at each iteration of the registration using known correspondence of gene markers and their associations with the AllenCCFv3. As previously reported,^65^ further assessment of the alignment showed that, of the 554 terminal regions (gray matter only) in the AllenCCFv3, only seven small subregions were missed from the MERFISH dataset: frontal pole, layer 1 (FRP1), FRP2/3, FRP5; accessory olfactory bulb, glomerular layer (AOBgl); accessory olfactory bulb, granular layer (AOBgr); accessory olfactory bulb, mitral layer (AOBmi); and accessory supraoptic group (ASO).

### 2.2 The DevCCF velocity flow model

**Figure 2:**
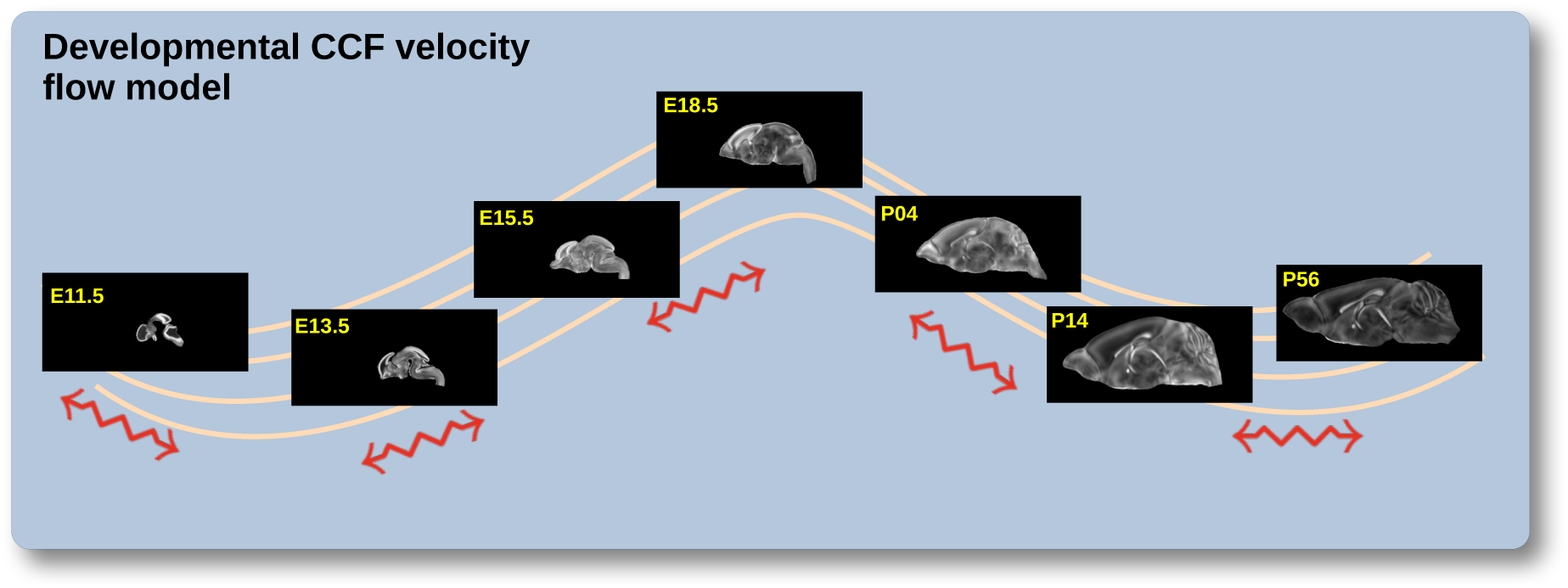
The spatial transformation between any two time points within the DevCCF longitudinal developmental trajectory is available through the use of ANTsX functionality for generating a velocity flow model.

To continuously interpolate transformations between the different stages of the DevCCF atlases, a velocity flow model was constructed using DevCCF derived data and functionality recently introduced into both the ANTsR and ANTsPy packages. Both platforms include a complete suite of functions for determining dense correspondence from sparse landmarks based on a variety of transformation models ranging from standard linear models (i.e., rigid, affine) to deformable diffeomorphic models (e.g, symmetric normalization).^42^ The latter set includes transformation models for both the pairwise scenario and for multiple sets, as in the case of the DevCCF. ANTsX, being built on top of ITK, uses an ITK image data structure for the 4-D velocity field where each voxel contains the *x*, *y*, *z* components of the field at that point.

#### 2.2.1 Data

Labeled annotations are available as part of the original DevCCF and reside in the space of each developmental template which range in resolution from 31.5 *−* 50*µ*m. Across all atlases, the total number of labeled regions exceeds 2500. From these labels, a common set of 26 labels (13 per hemisphere) across all atlases were used for optimization and evaluation. These simplified regions include: terminal hypothalamus, subpallium, pallium, peduncular hypothalamus, prosomere, prosomere, prosomere, midbrain, prepontine hindbrain, pontine hindbrain, pontomedullary hindbrain, medullary hindbrain, and tracts (see Figure 3).

**Figure 3:**
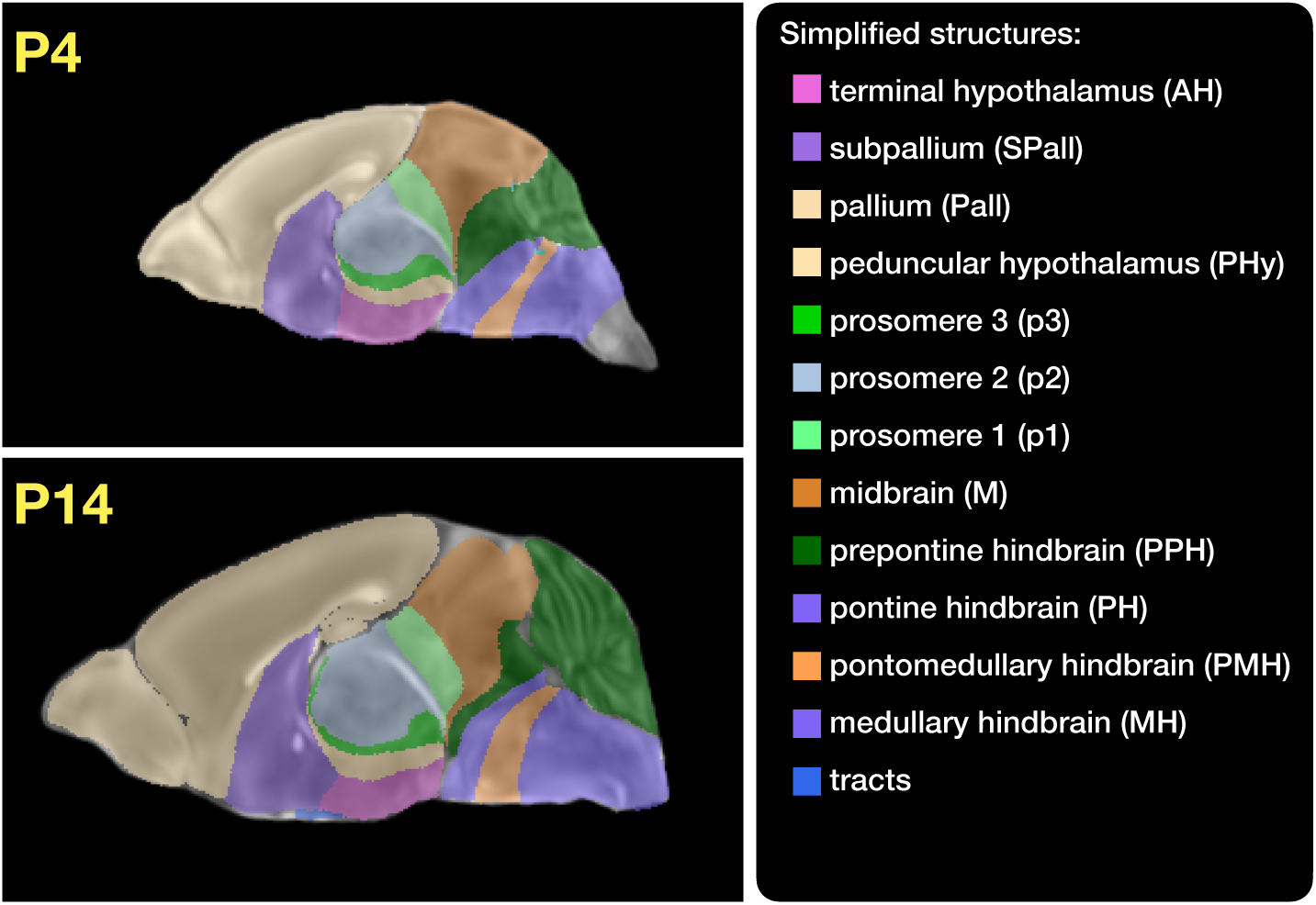
Annotated regions representing common labels across developmental stages which are illustrated for both P4 and P14.

Prior to velocity field optimization, all data were rigidly transformed to DevCCF P56 using the centroids of the common label sets. In order to determine the landmark correspondence across DevCCF stages, the multi-metric capabilities of ants.registration(…) were used. Instead of performing intensity-based pairwise registration directly on these multi-label images, each label was used to construct a separate fixed and moving image pair resulting in a multi-metric registration optimization scenario involving 24 binary image pairs (each label weighted equally) for optimizing diffeomorphic correspondence between neighboring time point atlases using the mean squares metric and the symmetric normalization transform.^42^

To generate the set of common point sets across all seven developmental atlases, the label boundaries and whole regions were sampled in the P56 atlas and then propagated to each atlas using the transformations derived from the pairwise registrations. We selected a sampling rate of 10% for the contour points and 1% for the regional points for a total number of points being per atlas being 173303 (*N_contour_*= 98151 and *N_region_*= 75152). Regional boundary points were weighted twice as those of non-boundary points during optimization.

#### 2.2.2 Optimization

The velocity field was optimized using the input composed of the seven corresponding point sets and their associated weight values, the selected number of integration points for the velocity field (*N* = 11), and the parameters defining the geometry of the spatial dimensions of the velocity field. Thus, the optimized velocity field described here is of size [256, 182, 360] (50*µ*m isotropic) ×11 integration points for a total compressed size of a little over 2 GB. This choice represented weighing the trade-off between tractability, portability, and accuracy. However, all data and code to reproduce the results described (with possible variation in the input parameters) are available in the dedicated GitHub repository.

The normalized time point scalar value for each atlas/point-set in the temporal domains [0, 1] was also defined. Given the increasingly larger gaps in the postnatal timepoint sampling, we made two adjustments. Based on known mouse brain development, we used 28 days for the P56 data. We then computed the log transform of the adjusted set of time points prior to normalization between 0 and 1 (see the right side of Figure 4). This log transform, as part of the temporal normalization, significantly improved data spacing.

**Figure 4:**
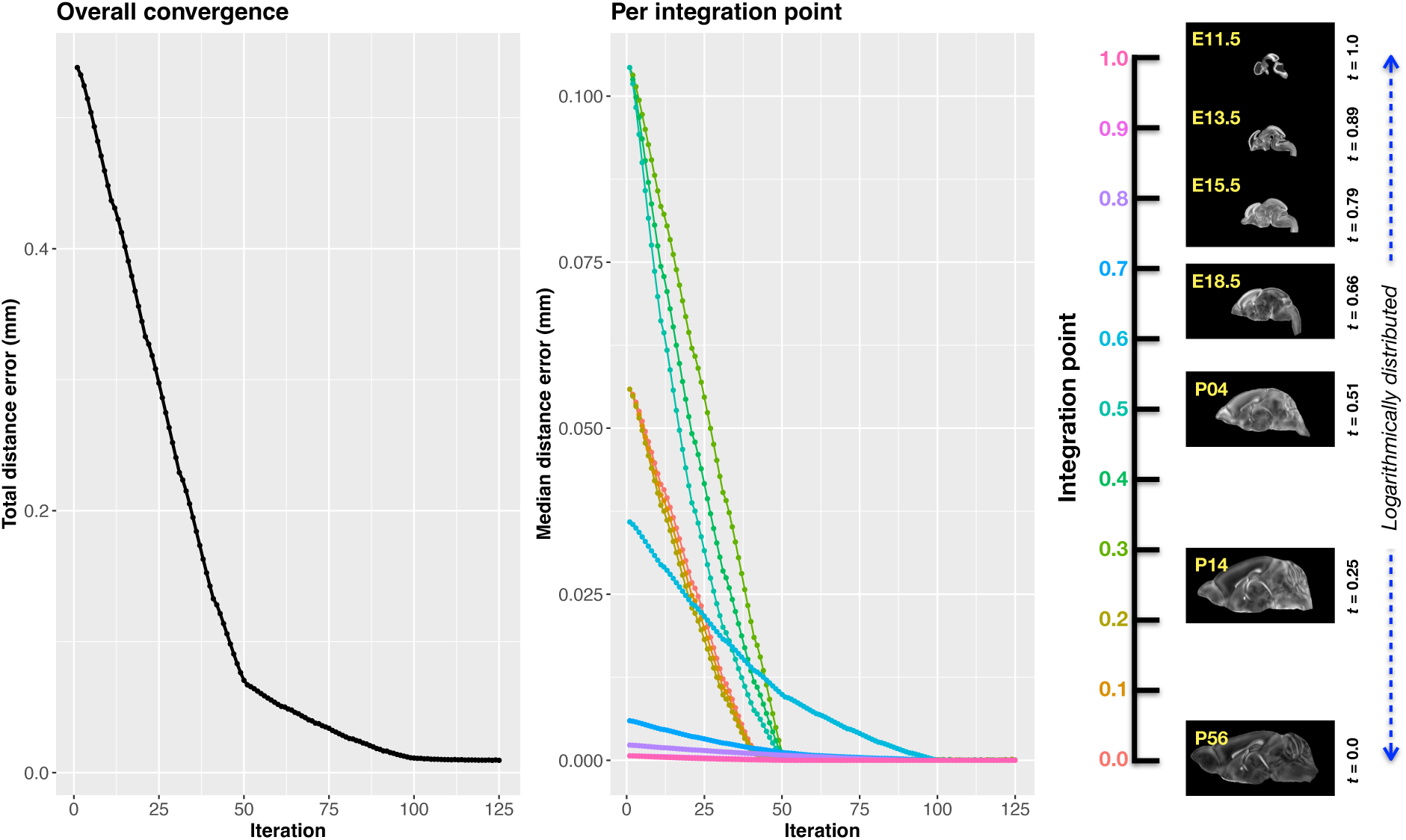
Convergence of the optimization of the velocity field for describing the transformation through the developmental stages from E11.5 through P56. Integration points in diagram on the right are color-coordinated with the center plot and placed in relation to the logarithmically situated temporal placement of the individual DevCCF atlases.

The maximum number of iterations was set to 200 with each iteration taking approximately six minutes on a 2020 iMac (processor, 3.6 GHz 10-Core Intel Core i9; memory, 64 GB 2667 MHz DDR4) At each iteration we looped over the 11 integration points. At each integration point, the velocity field estimate was updated by warping the two immediately adjacent point sets to the integration time point and determining the regularized displacement field between the two warped point sets. As with any gradient-based descent algorithm, this field was multiplied by a small step size (*δ* = 0.2) before adding to the current velocity field. Convergence is determined by the average displacement error over each of the integration points. As can be seen in the left panel of Figure 4, convergence occurred around 125 iterations when the average displacement error over all integration points is minimized. The median displacement error at each of the integration points also trends towards zero but at different rates.

#### 2.2.3 The transformation model

Once optimized, the resulting velocity field can be used to generate the deformable transform between any two continuous points within the time interval bounded by E11.5 and P56. In Figure 5, we transform each atlas to the space of every other atlas using the DevCCF transform model. Additionally, one can use this transformation model to construct virtual templates in the temporal gaps of the DevCCF. Given an arbitrarily chosen time point within the normalized time point interval, the existing adjacent DevCCF atlases on either chronological side can be warped to the desired time point. A subsequent call to one of the ANTsX template building functions then permits the construction of the template at that time point. In Figure 6, we illustrate the use of the DevCCF velocity flow model for generating two such virtual templates for two arbitrary time points. Note that both of these usage examples can be found in the GitHub repository previously given.

**Figure 5:**
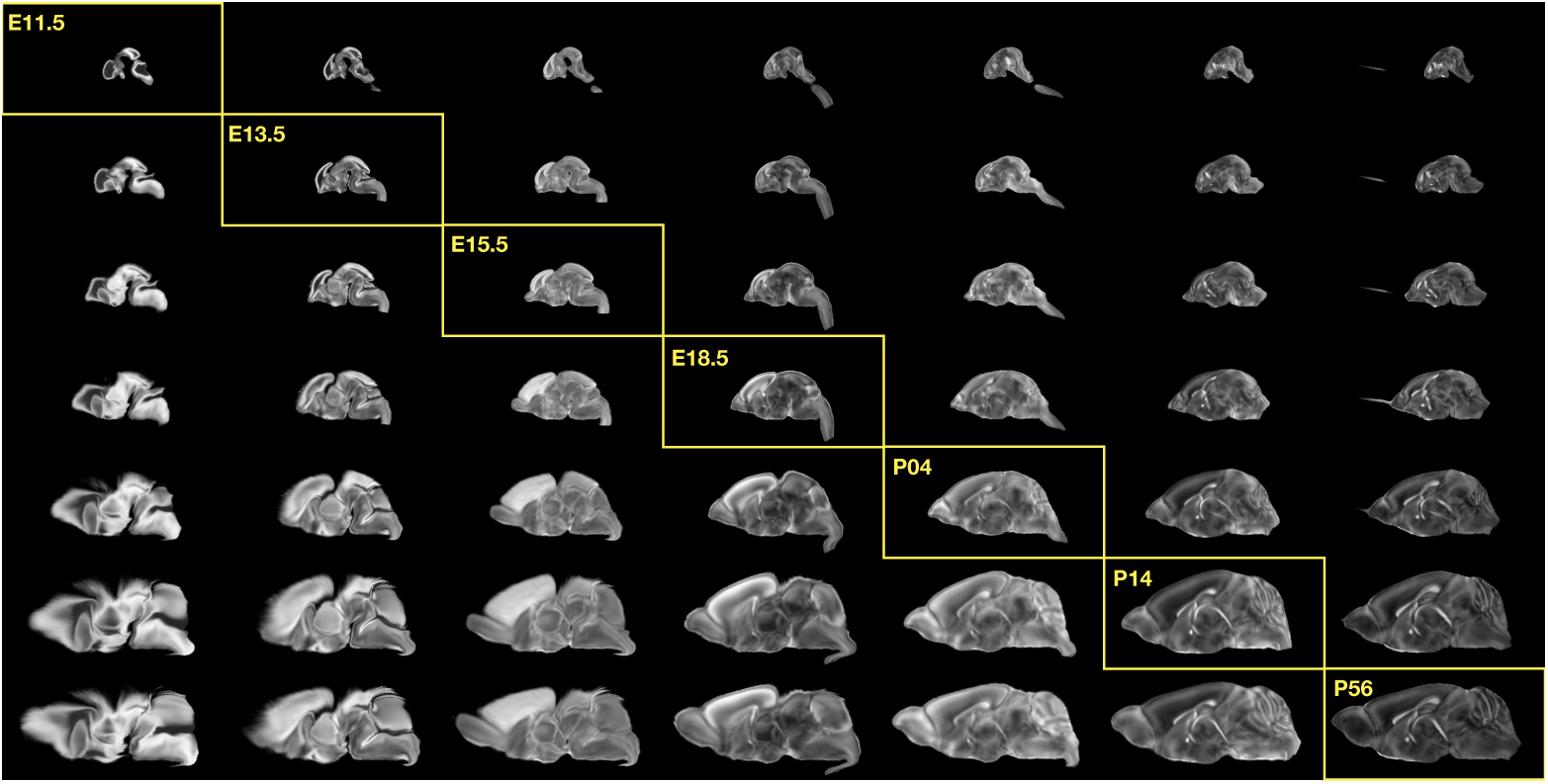
Mid-sagittal visualization of the effects of the transformation model in warping every developmental stage to the time point of every other developmental stage. The original images are located along the diagonal. Columns correspond to the warped original image whereas the rows represent the reference space to which each image is warped.

**Figure 6:**
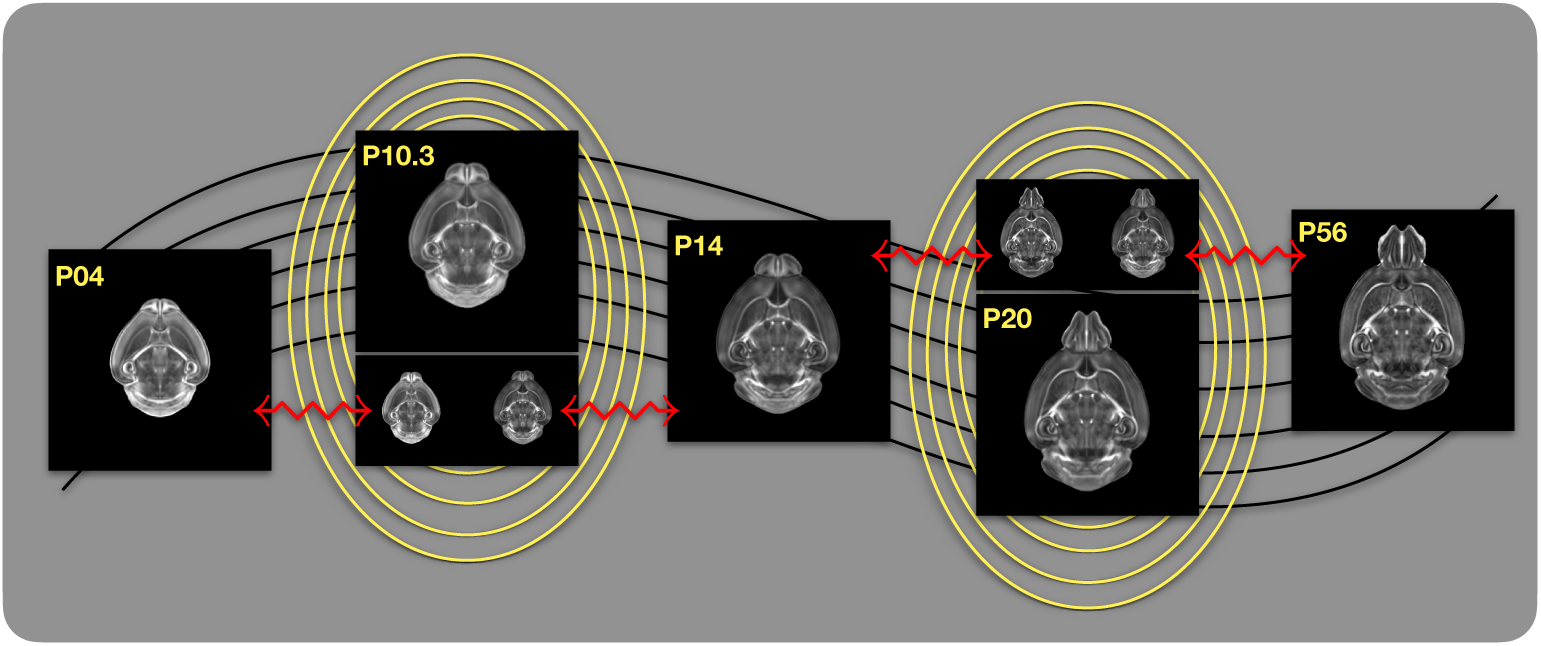
Illustration of the use of the velocity flow model for creating virtual templates at continuous time points not represented in one of the existing DevCCF time points. For example, FA templates at time point P10.3 and P20 can be generated by warping the existing temporally adjacent developmental templates to the target time point and using those images in the ANTsX template building process.

### 2.3 The Mouse Cortical Thickness Pipeline

One of the most well-utilized pipelines in the ANTsX toolkit is the generation of cortical thickness maps in the human brain from T1-weighted MRI. Starting with the novel Diffeomorphic Registration-based Cortical Thickness (DiReCT) algorithm,^55^ a complete algorithmic workflow was developed for both cross-sectional^56^ and longitudinal^57^ T1-weighted MR image data. This contribution was later refactored using deep learning^26^ leveraging the earlier results^56^ for training data.

In the case of the mouse brain, the lack of training data and/or tools to generate training data making analogous algorithmic development difficult. In addition, mouse data is often characterized by unique issues such as frequent anisotropic sampling which are often in sharp contrast to the high resolution resources available within the community, e.g., AllenCCFv3 and DevCCF. Using ANTsX and other publicly available data resources, we developed a complete mouse brain structural morphology pipeline as illustrated in Figure 7 and detailed below.

**Figure 7:**
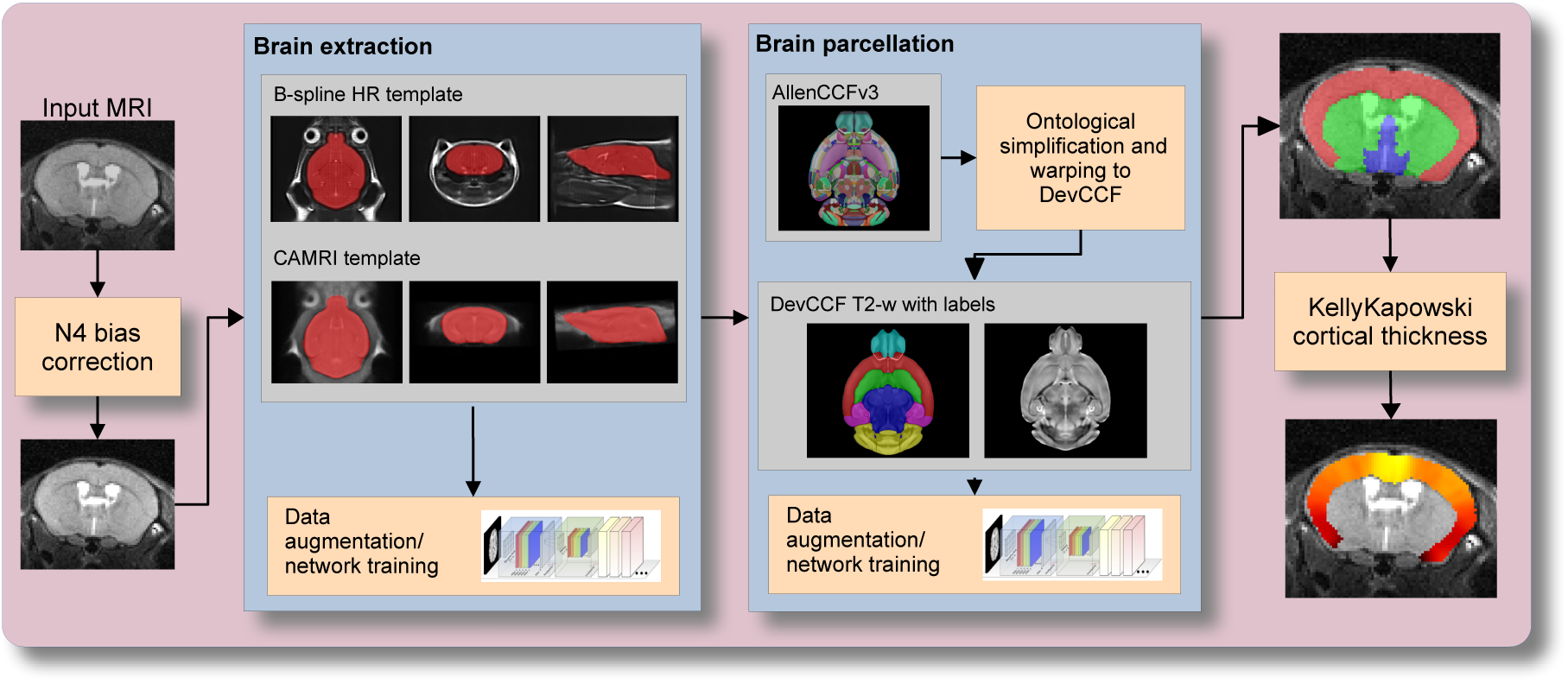
The mouse brain cortical thickness pipeline integrating two deep learning components for brain extraction and brain parcellation prior to estimating cortical thickness. Both deep learning networks rely heavily on data augmentation on templates built from open data and provide an outline for further refinement and creating alternative parcellations for tailored research objectives.

#### 2.3.1 Two-shot mouse brain extraction network

In order to create a generalized mouse brain extraction network, we built whole-head templates from two publicly available datasets. The Center for Animal MRI (CAMRI) dataset^58^ from the University of North Carolina at Chapel Hill consists of 16 T2-weighted MRI volumes of voxel resolution 0.16 × 0.16 × 0.16*mm*^3^. The second high-resolution dataset^59^ comprises 88 specimens each with three spatially aligned canonical views with in-plane resolution of 0.08 × 0.08*mm*^2^ with a slice thickness of 0.5*mm*. These three orthogonal views were used to reconstruct a single high-resolution volume per subject using a B-spline fitting algorithm developed in ANTsX.^68^ From these two datasets, two symmetric isotropic ANTsX templates^46^ were generated analogous to the publicly available ANTsX human brain templates used in previous research.^56^ Bias field simulation, intensity histogram warping, noise simulation, random translation and warping, and random anisotropic resampling in the three canonical directions were used for data augmentation in training a T2-weighted brain extraction network.

#### 2.3.2 Single-shot mouse brain parcellation network

To create the network for generating a brain parcellation consistent with cortical thickness estimation, we used the AllenCCFv3 and the associated allensdk Python library. Using allensdk, a gross parcellation labeling was generated from the fine Allen CCFv3 labeling which includes the cerebral cortex, cerebral nuclei, brain stem, cerebellum, main olfactory bulb, and hippocampal formation. This labeling was mapped to the P56 component of the DevCCF. Both the T2-w P56 DevCCF and labelings, in conjunction with the data augmentation described previously for brain extraction, was used to train a brain parcellation network.

#### 2.3.3 Evaluation

For evaluation, we used an additional publicly available dataset^60^ which is completely independent from the data used in training the brain extraction and parcellation networks. Data includes 12 specimens each imaged at seven time points (Day 0, Day 3, Week 1, Week 4, Week 8, Week 20) with available brain masks. In-plane resolution is 0.1 × 0.1*mm*^2^ with a slice thickness of 0.5*mm*. Since the training data is isotropic and data augmentation includes downsampling in the canonical directions, each of the two networks learns mouse brain-specific interpolation such that one can perform prediction on thick-sliced images, as, for example, in these evaluation data, and return isotropic probability and thickness maps (a choice available to the user). Figure 8 summarizes the results of the evaluation and comparison between isotropic and anisotropic cortical measurements in male and female specimens.

**Figure 8:**
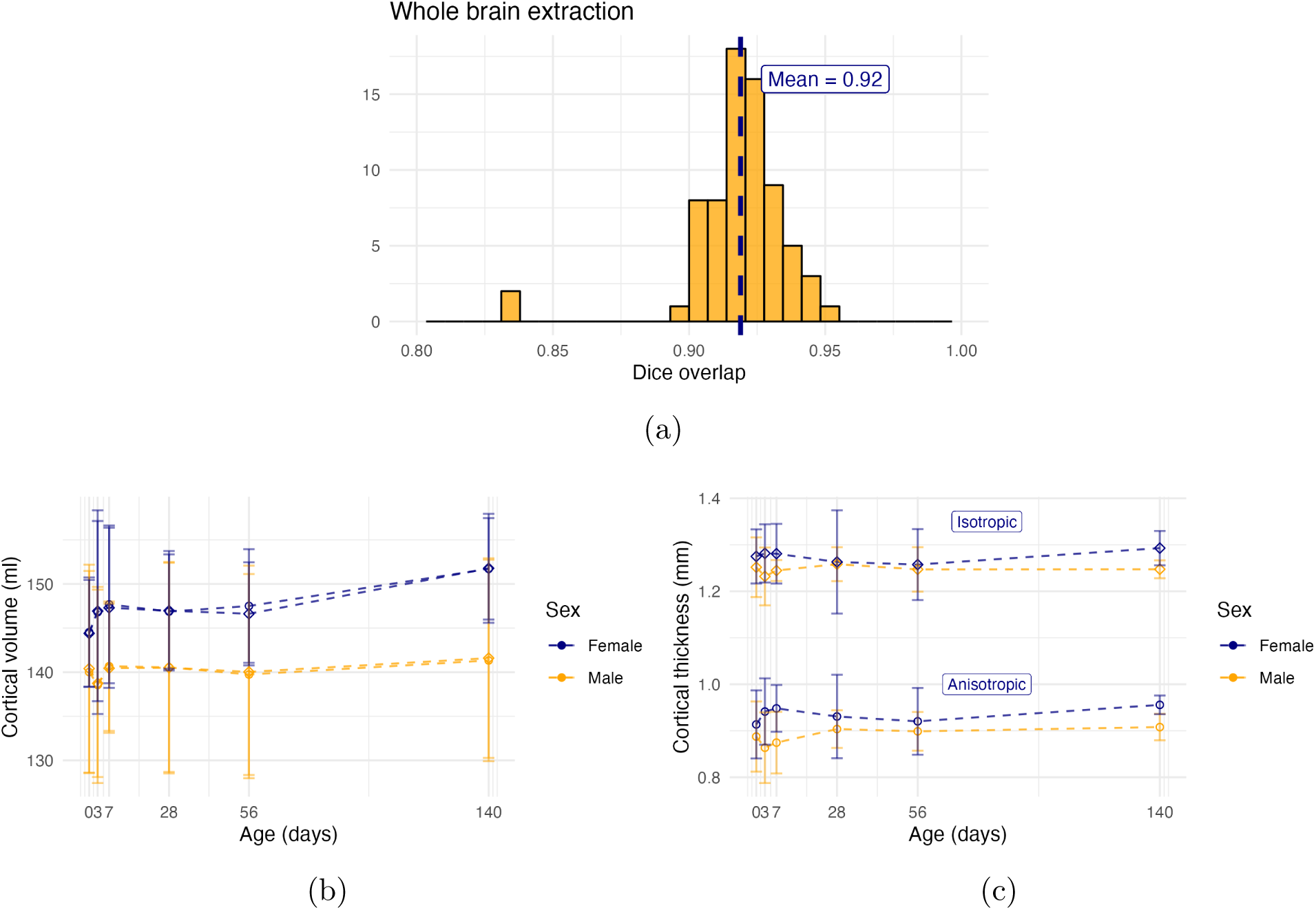
Evaluation of the ANTsX mouse brain extraction, parcellation, and cortical thickness pipeline on an independent dataset consisting of 12 specimens × 7 time points = 84 total images. (a) Dice overlap comparisons with the provided brain masks provide generally good agreement with the brain extraction network. (b) Cortical volume measurements show similar average quantities over growth and development between the original anisotropic data and interpolated isotropic data. (c) These results contrast with the cortical thickness measurements which show that cortical thickness estimation in anisotropic space severely underestimates the actual values.

## 3 Discussion

The ANTsX ecosystem is a powerful framework that has demonstrated applicability to multiple species and organ systems, including the mouse brain. This is further evidenced by the many software packages that use various ANTsX components in their own mouse-specific workflows. In and of itself, the extensive functionality of ANTsX makes it possible to create complete processing pipelines without requiring the integration of multiple packages. These open-source components not only perform well but are available across multiple platforms which facilitates the construction of tailored pipelines for individual study solutions. These components are also supported by years of development not only by the ANTsX development team but by the larger ITK community.

In the case of the development of the DevCCF, ANTsX was crucial in providing necessary functionality for yielding high quality output. For the generation of the individual developmental stage multi-modal, symmetric templates, ANTsX is unique amongst image analysis software packages in providing existing solutions for template generation which have been thoroughly vetted, including being used in several studies over the years, and which continue to be under active refinement. At its core, computationally efficient and quality template generation requires the use of precision pairwise image mapping functionality which, historically, is at the origins of the ANTsX ecosystem. Moreover, these mapping capabilities extend beyond template generation to the mapping of other image data (e.g., gene expression maps) to a selected template for providing further insight into the mouse brain.

With respect to the DevCCF, despite the significant expansion of available developmental age templates beyond what existed previously, there are still temporal gaps in the DevCCF which can be potentially sampled by future research efforts. However, pioneering work involving time-varying diffeomorphic transformations allow us to continuously situate the existing templates within a velocity flow model. This allows one to determine the diffeomorphic transformation from any one temporal location to any other temporal location within the time span defined by the temporal limits of the DevCCF. This functionality is built on multiple ITK components including the B-spline scattered data approximation technique for field regularization and velocity field integration. This velocity field model permits intra-template comparison and the construction of virtual templates where a template can be estimated at any continuous time point within the temporal domain. This novel application can potentially enhance our understanding of intermediate developmental stages.

We also presented a mouse brain pipeline for brain extraction, parcellation, and cortical thickness using single-shot and two-shot learning with data augmentation. This approach attempts to circumvent (or at least minimize) the typical requirement of large training datasets as with the human ANTsX pipeline analog. However, even given our initial success on independent data, we fully anticipate that refinements will be necessary. Given that the ANTsX toolkit is a dynamic effort undergoing continual improvement, we manually correct cases that fail and use them for future training and refinement of network weights as we have done for our human-based networks. Generally, these approaches provide a way to bootstrap training data for manual refinement and future generation of more accurate deep learning networks in the absence of other applicable tools.

## 4 Methods

The following methods are all available as part of the ANTsX ecosystem with analogous elements existing in both ANTsR (ANTs in R) and ANTsPy (ANTs in Python) with an ANTs/ITK C++ core. However, most of the development for the work described below was performed using ANTsPy. For equivalent calls in ANTsR, please see the ANTsX tutorial at https://tinyurl.com/antsxtutorial.

### 4.1 General ANTsX utilities

#### 4.1.1 Preprocessing: bias field correction and denoising

Bias field correction and image denoising are standard preprocessing steps in improving overall image quality in mouse brain images. The bias field, a gradual spatial intensity variation in images, can arise from various sources such as magnetic field inhomogeneity or acquisition artifacts, leading to distortions that can compromise the quality of brain images. Correcting for bias fields ensures a more uniform and consistent representation of brain structures, enabling more accurate quantitative analysis. Additionally, brain images are often susceptible to various forms of noise, which can obscure subtle features and affect the precision of measurements. Denoising techniques help mitigate the impact of noise, enhancing the signal-to-noise ratio and improving the overall image quality. The well-known N4 bias field correction algorithm^25^ has its origins in the ANTs toolkit which was implemented and introduced into the ITK toolkit, i.e. ants.n4_bias_field_correction(…). Similarly, ANTsX contains an implementation of a well-performing patch-based denoising technique^48^ and is also available as an image filter to the ITK community, ants.denoise_image(…).

#### 4.1.2 Image registration

The ANTs registration toolkit is a complex framework permitting highly tailored solutions to pairwise image registration scenarios.^69^ It includes innovative transformation models for biological modeling^42,54^ and has proven capable of excellent performance.^43,70^ Various parameter sets targeting specific applications have been packaged with the different ANTsX packages, specifically ANTs, ANTsPy, and ANTsR.^26^ In ANTsPy, the function ants.registration(…) is used to register a pair of images or a pair of image sets where type_of_transform is a user-specified option that invokes a specific parameter set. For example type_of_transform=`’`antsRegistrationSyNQuick[s]`’` encapsulates an oft-used parameter set for quick registration whereas type_of_transform=`’`antsRegistrationSyN[s]`’` is a more detailed alternative. Transforming images using the derived transforms is performed via the ants.apply_transforms(…) function.

Initially, linear optimization is initialized with center of (intensity) mass alignment typically followed by optimization of both rigid and affine transforms using the mutual information similarity metric. This is followed by diffeomorphic deformable alignment using symmetric normalization (SyN) with Gaussian^42^ or B-spline regularization^54^ where the forward transform is invertible and differentiable. The similarity metric employed at this latter stage is typically either neighborhood cross-correlation or mutual information. Note that these parameter sets are robust to input image type (e.g., light sheet fluorescence microscopy, Nissl staining, and the various MRI modalities) and are adaptable to mouse image geometry and scaling. Further details can be found in the various documentation sources for these ANTsX packages.

#### 4.1.3 Template generation

ANTsX provides functionality for constructing templates from a set (or multi-modal sets) of input images as originally described^46^ and recently used to create the DevCCF templates.^15^ An initial template estimate is constructed from an existing subject image or a voxelwise average derived from a rigid pre-alignment of the image population. Pairwise registration between each subject and the current template estimate is performed using the Symmetric Normalization (SyN) algorithm.^42^ The template estimate is updated by warping all subjects to the space of the template, performing a voxelwise average, and then performing a “shape update” of this latter image by warping it by the average inverse deformation, thus yielding a mean image of the population in terms of both intensity and shape. The corresponding ANTsPy function is ants.build_template(…).

#### 4.1.4 Visualization

To complement the well-known visualization capabilities of R and Python, e.g., ggplot2 and matplotlib, respectively, image-specific visualization capabilities are available in the ants.plot(…) function (Python). These are capable of illustrating multiple slices in different orientations with other image overlays and label images.

### 4.2 Mapping fMOST data to AllenCCFv3

#### 4.2.1 Preprocessing

- *Downsampling*. The first challenge when mapping fMOST images into the AllenCCFv3 is addressing the resolution scale of the data. Native fMOST data from an individual specimen can range in the order of terabytes, which leads to two main problems. First, volumetric registration methods (particularly those estimating local deformation) have high computational complexity and typically cannot operate on such high-resolution data under reasonable memory and runtime constraints. Second, the resolution of the AllenCCFv3 atlas is much lower than the fMOST data, thus the mapping process will cause much of the high-resolution information in the fMOST images to be lost regardless. Thus, we perform a cubic B-spline downsampling of the fMOST data to reduce the resolution of each image to match the isotropic 25 *µm* voxel resolution of the AllenCCFv3 intensity atlas using ants.resample_image(…). An important detail to note is that while the fMOST images and atlas are downsampled, the mapping learned during the registration is assumed to be continuous. Thus, after establishing the mapping to the AllenCCFv3, we can interpolate the learned mapping and apply it directly to the high-resolution native data directly to transform any spatially aligned data (such as the single-cell neuron reconstructions) into the AllenCCFv3.
- *Stripe artifact removal*. Repetitive pattern artifacts are a common challenge in fMOST imaging where inhomogeneity during the cutting and imaging of different sections can leave stripes of hyper- and hypo-intensity across the image. These stripe artifacts can be latched onto by the registration algorithm as unintended features that are then misregistered to non-analogous structures in the AllenCCFv3. We address these artifacts by fitting a 3-D bandstop (notch) filter to target the frequency of the stripe patterns and removing them prior to the image registration.
- *Inhomogeneity correction*. Regional intensity inhomogeneity can also occur within and between sections in fMOST imaging due to staining or lighting irregularity during acquisition. Similar to stripe artifacts, intensity gradients due to inhomogeneity can be misconstrued as features during the mapping and result in matching of non-corresponding structures. Our pipeline addresses these intensity inhomogeneities using N4 bias field correction,^25^ ants.n4_bias_field_correction(…).

#### 4.2.2 Steps for spatial normalization to AllenCCFv3

1. *Average fMOST atlas as an intermediate target*. Due to the preparation of the mouse brain for fMOST imaging, the resulting structure in the mouse brain has several large morphological deviations from the AllenCCFv3 atlas. Most notable of these is an enlargement of the ventricles, and compression of cortical structures. In addition, there is poor intensity correspondence for the same anatomic features due to intensity dissimilarity between imaging modalities. We have found that standard intensity-base registration is insufficient to capture the significant deformations required to map these structures correctly into the AllenCCFv3. We address this challenge in ANTsX by using explicitly corresponding parcellations of the brain, ventricles and surrounding structures to directly recover these large morphological differences. However, generating these parcellations for each individual mouse brain is a labor-intensive task. Our solution is to create an average atlas whose mapping to AllenCCFv3 encapsulates these large morphological differences to serve as an intermediate registration point. This has the advantage of only needing to generate one set of corresponding annotations which is used to register between the two atlas spaces. New images are first aligned to the fMOST average atlas, which shares common intensity and morphological features and thus can be achieved through standard intensity-based registration.
2. *Average fMOST atlas construction*. An intensity and shape-based contralaterally symmetric average of the fMOST image data is constructed from 30 images and their contralateral flipped versions. We ran three iterations of the atlas construction using the default settings. Additional iterations (up to six) were evaluated and showed minimal changes to the final atlas construction, suggesting a convergence of the algorithm.
3. *fMOST atlas to AllenCCFv3 alignment*. Alignment between the fMOST average atlas and AllenCCFv3 was performed using a one-time annotation-driven approach. Label-to-label registration is used to align 7 corresponding annotations in both atlases in the following: 1) brain mask/ventricles, 2) caudate/putamen, 3) fimbria, 4) posterior choroid plexus, 5) optic chiasm, 6) anterior choroid plexus, and 7) habenular commissure. The alignments were performed sequentially, with the largest, most relevant structures being aligned first using coarse registration parameters, followed by other structures using finer parameters. This coarse-to-fine approach allows us to address large morphological differences (such as brain shape and ventricle expansion) at the start of registration and then progressively refine the mapping using the smaller structures. The overall ordering of these structures was determined manually by an expert anatomist, where anatomical misregistration after each step of the registration was evaluated and used to determine which structure should be used in the subsequent iteration to best improve the alignment. The transformation from this one-time expert-guided alignment is preserved and used as the canonical fMOST atlas to AllenCCFv3 mapping in the pipeline.
4. *Alignment of individual fMOST mouse brains*. The canonical transformation between the fMOST atlas and AllenCCFv3 greatly simplifies the registration of new individual fMOST mouse brains into the AllenCCFv3. Each new image is first registered into the fMOST average atlas, which shares intensity, modality, and morphological characteristics. This allows us to leverage standard, intensity-based registration functionality^69^ available in ANTsX to perform this alignment. Transformations are then concatenated to the original fMOST image to move it into the AllenCCFv3 space using ants.apply_transforms(…).
5. *Transformation of single cell neurons*. A key feature of fMOST imaging is the ability to reconstruct and examine whole-brain single neuron projections^64^. Spatial mapping of these neurons from individual brains into the AllenCCFv3 allows investigators to study different neuron types within the same space and characterize their morphology with respect to their transcriptomics. Mappings found between the fMOST image and the AllenCCFv3 using our pipeline can be applied in this way to fMOST neuron reconstruction data.

### 4.3 Mapping MERFISH data to AllenCCFv3

#### 4.3.1 Preprocessing

- *Initial volume reconstruction*. Alignment of MERFISH data into a 3-D atlas space requires an estimation of anatomical structure within the data. For each section, this anatomic reference image was created by aggregating the number of detected genetic markers (across all probes) within each pixel of a 10 × 10*µm*^2^ grid to match the resolution of the 10*µm* AllenCCFv3 atlas. These reference image sections are then coarsely reoriented and aligned across sections using manual annotations of the most dorsal and ventral points of the midline. The procedure produces an anatomic image stack that serves as an initialization for further global mappings into the AllenCCFv3.
- *Anatomical correspondence labeling*. Mapping the MERFISH data into the AllenCCFv3 requires us to establish correspondence between the anatomy depicted in the MERFISH and AllenCCFv3 data. Intensity-based features in MERFISH data are not sufficiently apparent to establish this correspondence, so we need to generate instead corresponding anatomical labelings of both images with which to drive registration. These labels are already available as part of the AllenCCFv3; thus, the main challenge is deriving analogous labels from the spatial transcriptomic maps of the MERFISH data. Toward this end, we assigned each cell from the scRNA-seq dataset to one of the following major regions: cerebellum, CTXsp, hindbrain, HPF, hypothalamus, isocortex, LSX, midbrain, OLF, PAL, sAMY, STRd, STRv, thalamus and hindbrain. A label map of each section was generated for each region by aggregating the cells assigned to that region within a 10 × 10*µm*^2^ grid. The same approach was used to generate more fine grained region specific landmarks (i.e., cortical layers, habenula, IC). Unlike the broad labels which cover large swaths of the section these regions are highly specific to certain parts of the section. Once cells in the MERFISH data are labeled, morphological dilation is used to provide full regional labels for alignment into the AllenCCFv3.
- *Section matching*. Since the MERFISH data is acquired as sections, its 3-D orientation may not be fully accounted for during the volume reconstruction step, due to the particular cutting angle. This can lead to obliqueness artifacts in the section where certain structures can appear to be larger or smaller, or missing outright from the section. To address this, we first use a global alignment to match the orientations of the MERFISH sections to the atlas space. In our pipeline, this section matching is performed in the reverse direction by performing a global affine transformation of the AllenCCFv3 into the MERFISH data space, and then resampling digital sections from the AllenCCFv3 to match each MERFISH section. This approach limits the overall transformation and thus resampling that is applied to the MERFISH data, and, since the AllenCCFv3 is densely sampled, it also reduces in-plane artifacts that result from missing sections or undefined spacing in the MERFISH data.

#### 4.3.2 2.5D deformable, landmark-driven alignment to AllenCCFv3

After global alignment of the AllenCCFv3 into the MERFISH dataset, 2D per-section deformable refinements are used to address local differences between the MERFISH sections and the resampled AllenCCFv3 sections. Nine registrations were performed in sequence using a single label at each iteration in the following order: 1) brain mask, 2) isocortex (layer 2+3), 3) isocortex (layer 5), 4) isocortex (layer 6), 5) striatum, 6) medial habenula, 7) lateral habenula, 8) thalamus, and 9) hippocampus. This ordering was determined empirically by an expert anatomist who prioritized which structure to use in each iteration by evaluating the anatomical alignment from the previous iteration. Global and local mappings are then all concatenated (with appropriate inversions) to create the final mapping between the MERFISH data and AllenCCFv3. This mapping is then used to provide a point-to-point correspondence between the original MERFISH coordinate space and the AllenCCFv3 space, thus allowing mapping of individual genes and cell types located in the MERFISH data to be directly mapped into the AllenCCFv3.

### 4.4 DevCCF velocity flow transformation model

Given multiple, linearly or non-linearly ordered point sets where individual points across the sets are in one-to-one correspondence, we developed an approach for generating a velocity flow transformation model to describe a time-varying diffeomorphic mapping as a variant of the landmark matching solution. Integration of the resulting velocity field can then be used to describe the displacement between any two time points within this time-parameterized domain. Regularization of the sparse correspondence between point sets is performed using a generalized B-spline scattered data approximation technique,^68^ also created by the ANTsX developers and contributed to ITK.

#### 4.4.1 Velocity field optimization

To apply this methodology to the developmental templates,^15^ we coalesced the manual annotations of the developmental templates into 26 common anatomical regions (see Figure 3). We then used these regions to generate invertible transformations between successive time points. Specifically each label was used to create a pair of single region images resulting in 26 pairs of “source” and “target” images. The multiple image pairs were simultaneously used to iteratively estimate a diffeomorphic pairwise transform. Given the seven atlases E11.5, E13.5, E15.5, E18.5, P4, P14, and P56, this resulted in 6 sets of transforms between successive time points. Approximately 10^6^ points were randomly sampled labelwise in the P56 template space and propagated to each successive atlas providing the point sets for constructing the velocity flow model. Approximately 125 iterations resulted in a steady convergence based on the average Euclidean norm between transformed point sets. Ten integration points were used and point sets were distributed along the temporal dimension using a log transform for a more evenly spaced sampling. For additional information a help menu is available for the ANTsPy function ants.fit_time_varying_transform_to_point_sets(…).

### 4.5 ANTsXNet mouse brain applications

#### 4.5.1 General notes regarding deep learning training

All network-based approaches described below were implemented and organized in the ANTsXNet libraries comprising Python (ANTsPyNet) and R (ANTsRNet) analogs using the Keras/Tensorflow libraries available as open-source in ANTsX GitHub repositories. For the various applications, both share the identically trained weights for mutual reproducibility. For all GPU training, we used Python scripts for creating custom batch generators which we maintain in a separate GitHub repository for public availability (https://github.com/ntustison/ANTsXNetTraining). These scripts provide details such as batch size, choice of loss function, and network parameters. In terms of GPU hardware, all training was done on a DGX (GPUs: 4X Tesla V100, system memory: 256 GB LRDIMM DDR4).

Data augmentation is crucial for generalizability and accuracy of the trained networks. Intensity-based data augmentation consisted of randomly added noise (i.e., Gaussian, shot, salt-and-pepper), simulated bias fields based on N4 bias field modeling, and histogram warping for mimicking well-known MRI intensity nonlinearities.^26,71^ These augmentation techniques are available in ANTsXNet (only ANTsPyNet versions are listed with ANTsRNet versions available) and include:

- image noise: ants.add_noise_to_image(…),
- simulated bias field: antspynet.simulate_bias_field(…), and
- nonlinear intensity warping: antspynet.histogram_warp_image_intensities(…).

Shape-based data augmentation used both random linear and nonlinear deformations in addition to anisotropic resampling in the three canonical orientations to mimic frequently used acquisition protocols for mice brains:

- random spatial warping: antspynet.randomly_transform_image_data(…) and
- anisotropic resampling: ants.resample_image(…).

#### 4.5.2 Brain extraction

Similar to human neuroimage processing, brain extraction is a crucial preprocessing step for accurate brain mapping. We developed similar functionality for T2-weighted mouse brains. This network uses a conventional U-net architecture^72^ and, in ANTsPyNet, this functionality is available in the program antspynet.mouse_brain_extraction(…). For the two-shot T2-weighted brain extraction network, two brain templates were generated along with their masks. One of the templates was generated from orthogonal multi-plane, high resolution data^59^ which were combined to synthesize isotropic volumetric data using the B-spline fitting algorithm.^68^ This algorithm is encapsulated in ants.fit_bspline_object_to_scattered_data(…) where the input is the set of voxel intensity values and each associated physical location. Since each point can be assigned a confidence weight, we use the normalized gradient value to more heavily weight edge regions. Although both template/mask pairs are available in the GitHub repository associated with this work, the synthesized volumetric B-spline T2-weighted pair is available within ANTsXNet through the calls:

- template: antspynet.get_antsxnet_data(“bsplineT2MouseTemplate”) and
- mask: antspynet.get_antsxnet_data(“bsplineT2MouseTemplateBrainMask”).

#### 4.5.3 Brain parcellation

The T2-weighted brain parcellation network is also based on a 3-D U-net architecture and the T2-w DevCCF P56 template component with extensive data augmentation, as described previously. Intensity differences between the template and any brain extracted input image are minimized through the use of the rank intensity transform (ants.rank_intensity(…)). Shape differences are reduced by the additional preprocessing step of warping the brain extracted input image to the template. Additional input channels include the prior probability images created from the template parcellation. These images are also available through the ANTsXNet get_antsxnet_data(…) interface.

## Data availability

All data and software used in this work are publicly available. The DevCCF atlas is available at https://kimlab.io/brain-map/DevCCF/. ANTsPy, ANTsR, ANTsPyNet, and ANTsRNet are available through GitHub at the ANTsX Ecosystem (https://github.com/ANTsX). Training scripts for all deep learning functionality in ANTsXNet can also be found on GitHub (https://github.com/ntustison/ANTsXNetTraining). A GitHub repository specifically pertaining to the AllenCCFv3 mapping is available at https://github.com/dontminchenit/CCFAlignmentToolkit. For the other two contributions contained in this work, the longitudinal DevCCF mapping and mouse cortical thickness pipeline, we refer the interested reader to https://github.com/ntustison/ANTsXMouseBrainMapping.

## Acknowledgments

Support for the research reported in this work includes funding from the National Institute of Biomedical Imaging and Bioengineering (R01-EB031722) and National Institute of Mental Health (RF1-MH124605 and U24-MH114827).

## Author contributions

N.T., M.C., and J.G. wrote the main manuscript text and figures. M.C., M.K., R.D., S.S., Q.W., L.G., J.D., C.G., and J.G. developed the Allen registration pipelines. N.T. and F.K. developed the time-varying velocity transformation model for the DevCCF. N.T. and M.T. developed the brain parcellation and cortical thickness methodology. All authors reviewed the manuscript.

